# Scaling up DNA data storage and random access retrieval

**DOI:** 10.1101/114553

**Authors:** Lee Organick, Siena Dumas Ang, Yuan-Jyue Chen, Randolph Lopez, Sergey Yekhanin, Konstantin Makarychev, Miklos Z. Racz, Govinda Kamath, Parikshit Gopalan, Bichlien Nguyen, Christopher Takahashi, Sharon Newman, Hsing-Yeh Parker, Cyrus Rashtchian, Kendall Stewart, Gagan Gupta, Robert Carlson, John Mulligan, Douglas Carmean, Georg Seelig, Luis Ceze, Karin Strauss

## Abstract

Current storage technologies can no longer keep pace with exponentially growing amounts of data. ^1^ Synthetic DNA offers an attractive alternative due to its potential information density of ~ 10^18^ B/mm^3^, 10^7^ times denser than magnetic tape, and potential durability of thousands of years.^2^ Recent advances in DNA data storage have highlighted technical challenges, in particular, coding and random access, but have stored only modest amounts of data in synthetic DNA. ^3,4,5^ This paper demonstrates an end-to-end approach toward the viability of DNA data storage with large-scale random access. We encoded and stored 35 distinct files, totaling 200MB of data, in more than 13 million DNA oligonucleotides (about 2 billion nucleotides in total) and fully recovered the data with no bit errors, representing an advance of almost an order of magnitude compared to prior work. ^6^ Our data curation focused on technologically advanced data types and historical relevance, including the Universal Declaration of Human Rights in over 100 languages,^7^ a high-definition music video of the band OK Go,^8^ and a CropTrust database of the seeds stored in the Svalbard Global Seed Vault.^9^ We developed a random access methodology based on selective amplification, for which we designed and validated a large library of primers, and successfully retrieved arbitrarily chosen items from a subset of our pool containing 10.3 million DNA sequences. Moreover, we developed a novel coding scheme that dramatically reduces the physical redundancy (sequencing read coverage) required for error-free decoding to a median of 5x, while maintaining levels of logical redundancy comparable to the best prior codes. We further stress-tested our coding approach by successfully decoding a file using the more error-prone nanopore-based sequencing. We provide a detailed analysis of errors in the process of writing, storing, and reading data from synthetic DNA at a large scale, which helps characterize DNA as a storage medium and justify our coding approach. Thus, we have demonstrated a significant improvement in data volume, random access, and encoding/decoding schemes that contribute to a whole-system vision for DNA data storage.

---

Storing digital data using synthetic DNA encompasses mapping bits into nucleotide sequences, synthesizing the corresponding molecules, and storing them in an appropriate environment. Reading the information requires sequencing and converting the stored DNA back into digital data. Our project explores this DNA data storage workflow end-to-end (**Figure *1a***). We focus on scaling up data volumes and solving associated key challenges. Specifically, we address the need to access data selectively rather than in bulk and the need to minimize the amount of sequencing required to completely recover stored data.

**Figure 1:**
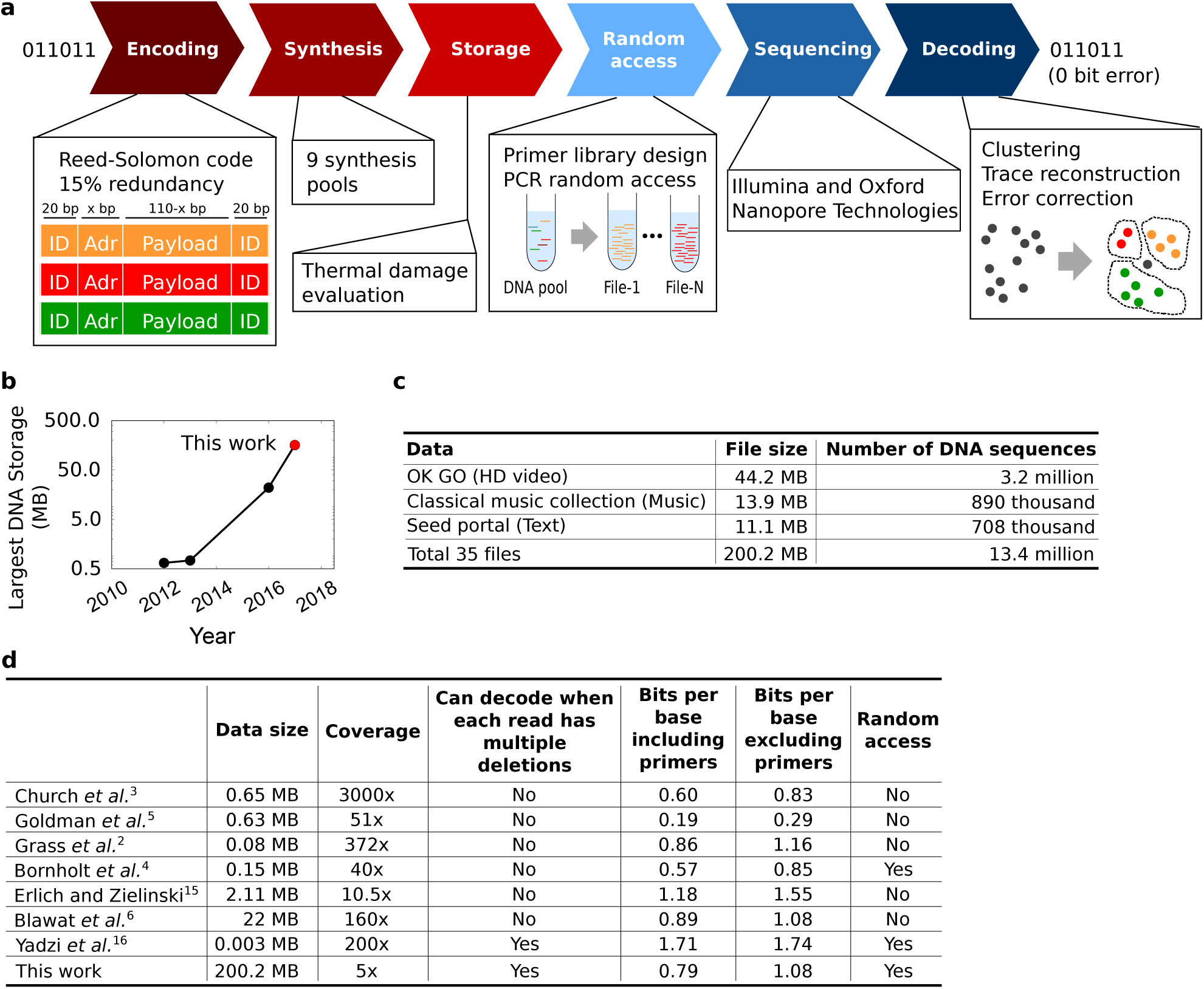
Overview of the DNA data storage workflow and stored data. **(a)** The encoding process maps digital files into a large set of 150-nucleotide DNA sequences, including Reed-Solomon code redundancy to overcome errors in synthesis and sequencing. The resulting collection of sequences is synthesized by Twist Bioscience. DNA pools are then stored under different conditions, including storage at high temperature meant to simulate the aging process. The random access process starts with amplifying a subset of the sequences corresponding to one of the files using polymerase chain reaction (PCR). The amplified pools are sequenced using either sequencing by synthesis (Illumina NextSeq) or nanopore sequencing (Oxford Nanopore Technologies). Finally, sequencing reads are decoded using our novel clustering and consensus algorithms. **(b)** We encoded a total of 200MB of data, about an order of magnitude more than prior work. **(c)** Example files encoded within these 200MB of data. **(d)** A comparison to prior work shows that our coding scheme has similar logical redundancy, but requires much lower physical redundancy and thus sequencing coverage, to recover files. Moreover, unlike most prior work we can select files using random access and decoding is possible even if all reads have multiple deletions.

Most prior DNA data storage research sequenced and decoded the entire amount of stored information, with no random access. However, such wholesale sequencing becomes impractical as the amount of data increases (**Figure *1b* and Figure *1c***). Being able to selectively access only part of the written information (e.g., retrieving only one image from a collection) is thus necessary to make DNA data storage viable, but so far has been demonstrated only at a small scale. ^4,10^ Our work demonstrates that PCR-based random access can be scaled up to reliably extract files of widely varying size and complexity from a DNA pool three orders of magnitude larger than those used in prior random access experiments. ^4^

Lack of random access is not the only issue limiting scale-up. Both DNA synthesis and sequencing are highly error prone processes resulting in aggregate insertion, deletion, and substitution rates observed at approximately 0.01 errors/base. ^11^ Even complete loss of specific data strands can occur during library synthesis or amplification. Prior work has shown that it is possible to recover data even from such noisy measurements if proper encoding schemes are used. Although efforts have been made to minimize the amount of *logical* redundancy (i.e., the amount of additional information encoded) required for complete data recovery at a given error rate, existing approaches rely on a high degree of *physical* redundancy, i.e., having many copies of each sequence and deep sequencing coverage. Here, we present a coding scheme that is unique in that it explicitly reduces physical redundancy, requiring fewer physical copies of any given molecule to fully recover the stored data. Our scheme tolerates aggressive settings of uneven low coverage and high coordinate error rates of insertions, deletions, and substitutions, while maintaining a logical density (bits per nucleotide) competitive with previously proposed schemes (**Figure *1d***). As DNA data storage technology matures, the goals of increasing throughput and lowering costs will likely drive coordinate error rates in the DNA data storage channel even higher than the current value.

To investigate challenges associated with increasing DNA data storage size, we created a large DNA library of modern data types, such as high definition video, images, audio, and text. We encoded 35 files ranging from 29 kilobytes to over 44 megabytes, totaling over 200MB of unique (compressed) data (**Figure *1c*** lists a few examples and **Supplementary Table *1*** provides the full list). We added 15% logical redundancy for robust error correction to 33 of our files and 25% to the other two, resulting in an additional 32.2MB of data encoded in DNA. For DNA synthesis, we segmented each input file into a large number of oligonucleotides, each containing the same PCR primer target sequences that form a unique file ID. Moreover, each strand also includes a unique, strand-specific address to order strands within a file. The resulting synthetic DNA library contains 13,448,372 unique DNA sequences of lengths ranging from 150 to 154 bases synthesized using Twist Bioscience’s oligo pool services in a total of 9 synthesis pools. Our resulting combined pool of about 2 billion bases represents an increase of about an order of magnitude in the amount of information stored in and retrieved from DNA, relative to prior work. ^6^

Achieving robust random access in a large DNA data storage system requires effective PCR primers to reliably amplify a specific file without crosstalk. We thus devised a framework for designing a primer library with thousands of pairs of orthogonal primers (file IDs). Our design method (**Figure *2a-i***) optimizes primers for several properties: avoidance of secondary structure formation and primer-dimer formation, absence of long stretches of homopolymers, melting temperature constrained to a narrow range (between 55°C and 60°C), and a minimum percentage of its sequence unique from other primers (30%). To increase the stringency of sequence orthogonality, we used the basic sequence alignment program BLAST to screen out primers with long stretches of similar sequences. ^12^ Before incorporating selected primer sequences into the library files, we tested primer performance on a pool of 3240 synthetic “mini-files” ranging from 1 to 200 103-mers. We successfully accessed and sequenced up to 48 mini-files from this pool in a one-pot multiplex PCR experiment (**Figure *2a-ii* and Figure *2a-iii***), validating our primer library design approach.

**Figure 2:**
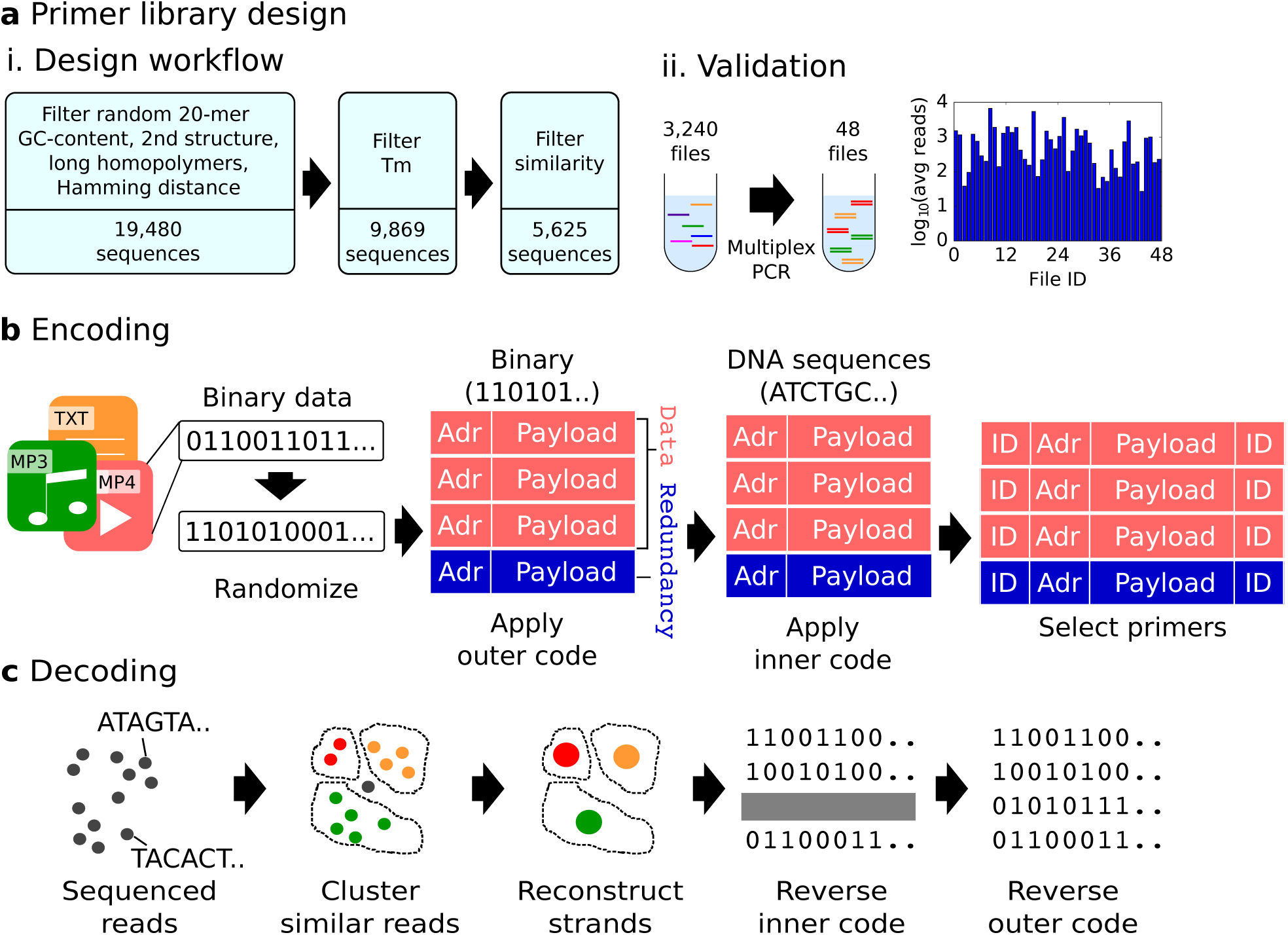
Design of random access primers and coding algorithm. **(a-i)** We designed a primer library for our PCR-based random access method using an in-silico process that starts with a random set of 20-mers. It filters them by melting temperature and then selects a set that is as diverse as possible, i.e., has low similarity between the sequences. **(a-ii)** The resulting set of candidate primers is then validated experimentally by synthesizing a pool of about 100 thousand strands containing sets of size 1 to 200 DNA sequences each, surrounded by one of the 3,240 candidate primer pairs, and then randomly selecting 48 of those pairs for amplification. The product is sequenced and sequences with each of the 48 primer pairs appear among sequencing reads, albeit at different relative proportions when normalized to the number of sequences in each set. **(b)** Our encoding process starts by randomizing data to reduce chances of secondary structures, primer-payload nonspecific binding (see Supplementary Figure 2), and improved properties during decoding. It then breaks the data into fixed-size payloads, adds addressing information, and applies outer coding, which adds redundant sequences using a Reed-Solomon code to increase robustness to missing sequences and errors. The level of redundancy is determined by expected error in sequencing and synthesis, as well as DNA degradation. Next, it applies inner coding, which ultimately converts the bits to DNA sequences. The resulting set of sequences is surrounded by a primer pair chosen from the library based on (low) level of overlap with payloads. **(c)** The decoding process starts by clustering reads based on similarity, and finding a consensus between the sequences in each cluster to reconstruct the original sequences, which are then decoded back to digital data.

Next, we created a coding scheme to convert digital information to DNA sequences and back to digital information. We begin by explaining our approach to data encoding (**Figure *2b***). Similar to prior work, ^2,6^ we employ concatenated codes with Reed-Solomon (RS) as the outer code.

However, unlike earlier work, we use very long codes (length 65,536) to handle large variations in the number of errors between codewords. Input data is randomized by performing an exclusive OR (XOR) operation with the output of a seeded pseudo-random number generator. Randomization facilitates coping with errors by breaking multi-bit repeats (e.g., 00000000) that pose a problem for decoding. The encoder then partitions the randomized digital file into RS codewords, each represented by sequences of bits that later will be translated into DNA sequences of length 110.

Next, the encoder adds redundant information for error correction and the bit sequences are converted into DNA sequences by using a rotating code that eliminates homopolymers. ^5^ As a result, when 15% redundancy is added, 87% of the DNA sequences will carry raw input data (systematic RS coordinates), while 13% will carry redundant data used for error correction (redundant RS coordinates). Every DNA sequence has the same format: a prefix with address information (its location in the file) followed by a payload (original data or redundant data). All DNA sequences are later appended with 20-base PCR primer targets selected from the primer library on both ends to allow random access to the file. Finally, the resulting DNA sequences are synthesized into DNA strands, which can then be preserved using a variety of methods, and later selected via random access.

Sequencing these DNA strands produces a collection of noisy reads, which do not necessarily include all original DNA sequences due to losses while sampling, moving, and operating over the DNA. Sequences belonging to a specific file are obtained by aligning and filtering based on the primer sequence and length. Frugality with respect to coverage was a key consideration when designing our decoding approach. As such, we do not require reads to be the correct length ^2^ and we do not filter out reads with errors. ^15^ Instead, noisy reads whose length is within 20 nucleotides of the original length are selected and passed to the decoder.

The decoder operates in four stages (**Figure *2c***). First, it clusters noisy reads by similarity based on their entire content, not just the addresses, ^6^ to collect all available reads that likely correspond to a unique originally stored DNA sequence. To do so, we employ an algorithm based on an iterative locality sensitive hashing designed for edit distance metric space, which relies on the input randomization done during encoding. The decoder then processes each cluster to recover the original sequence. This stage, which we call trace reconstruction, uses a variant of the Bitwise Majority Alignment algorithm ^13^ adapted to support insertions, deletions, and substitutions. In the third stage, the decoder unwinds the no-homopolymer representation to obtain indices and values of individual coordinates of the outer code. After stage three, some recovered sequences may be missing (erasures), and others may be incorrect (errors). In stage four, we decode the outer Reed-Solomon (RS) code to correct errors and erasures and invert randomization. Successful decoding is possible if for each RS codeword the ‘Utilized Error Resilience’ ratio 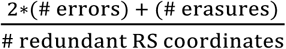 is at most 1.

We received the synthesized DNA pools periodically over several months. In each case, we immediately amplified every file in a pool using random access emulsion PCR, attached Illumina sequencing primers and adaptors, and then sequenced the files in a total of 10 sequencing runs. We have aggregated about 723 million reads of more than 13 million distinct synthetic DNA sequences. The mean coverage (i.e., number of reads for a given DNA sequence) across the dataset was 53.8 reads with a standard deviation of 48.7 reads. We observed considerable variance across files, ranging from a mean of 6.7 reads with standard deviation 3.4 reads to a mean of 298.6 reads with standard deviation of 139.6 reads. For most files, the empirical coverage distribution is reasonably well-approximated by a gamma distribution with matching mean and variance (**Figure *3a***, center).

**Figure 3:**
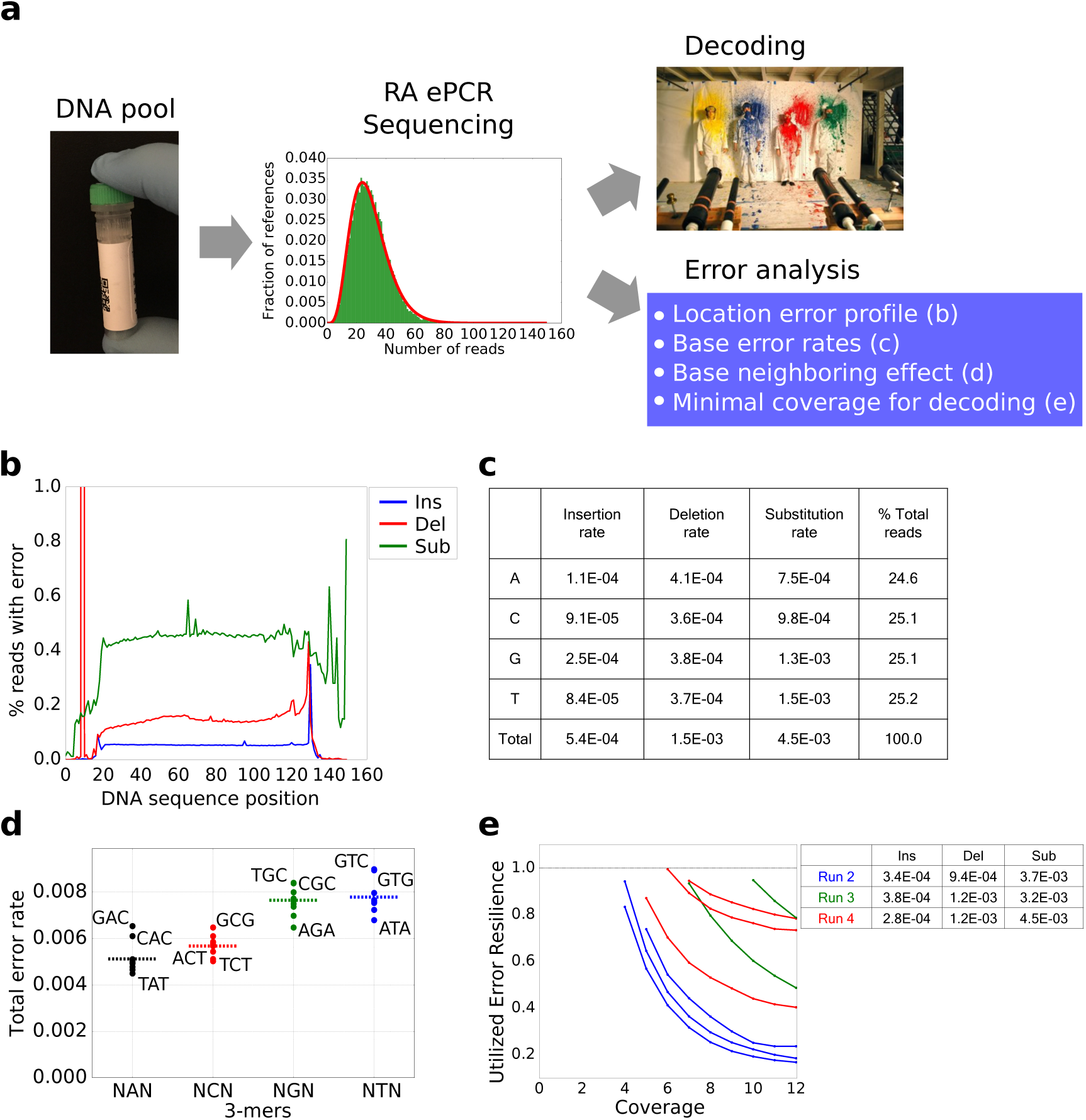
Experimental error analysis and decoding, sequencing using Illumina’s NextSeq. **(a)**Overview of the experimental analysis workflow: A file of interest is amplified from a stored pool and sequenced, resulting in a distribution of sequencing reads from which the file is decoded and the original stored data recovered, and a file-specific error profile is generated. **(b)** Per-position average read error profile averaged over the first 150 positions of all 13M strands and their corresponding reads. There is a spike to almost 15% at index 9 caused by an error in the primer region of a single file. **(c)** Error rates and number of reads for different nucleotide types in the payload region. Almost half of the insertions are associated to type G and about a third of the substitutions are associated to type T. Deletions are evenly distributed. **(d)** Error rates depend not only on nucleotide type but also on the type of neighboring nucleotides. Each dot corresponds to a 3-mer in the payload region colored according to the central base: black for A, red for C, green for G, and blue for T. Again, types G and T are associated with higher error rates. **(e)** Estimating the minimal coverage required for decoding. Each curve corresponds to a different file, each color corresponds to a different sequencing run, and numbers in the legend correspond to the average insertion, deletion, and substitution errors for the corresponding sequencing run. Redundant information is more scarce at lower coverages, resulting in higher utilized error resilience.

**Figure 4:**
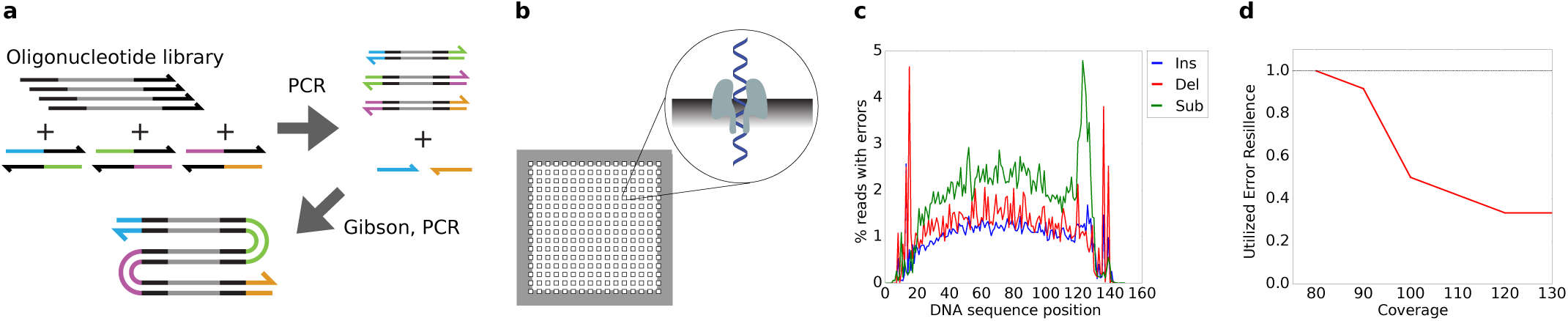
Sequencing using Oxford Nanopore Technologies' MinlON. Overview of DNA data storage workflow using Oxford Nanopore sequencing. **(a)** A file of interest is amplified using PCR with primers containing complementary overhang sequences. Subsequently, amplification products are mixed in a Gibson assembly reaction and amplified using primers corresponding to the unique overhangs present at the 5' and 3' of the Gibson assembly product. **(b)** Amplicon consisting of concatenated oligonucleotides is sequenced using the Oxford Nanopore platform and thousands of reads are generated. **(c)** Per-position average read error profile averaged over all 88 strands and their corresponding 2D reads. Error rates are higher than in Illumina-sequenced reads. **(d)** Estimating the minimal coverage required for decoding. Higher error rates are offset by higher coverage, making decoding the original stored data possible.

The sequencing information serves two purposes: (1) error analysis of processes related to DNA manipulation, including synthesis, random access, and sequencing, when used in conjunction with knowledge of the encoded DNA sequences; and (2) decoding of data stored in DNA and analysis of code resilience, i.e., its ability to recover the information under the observed error regime. The decoding process uses only information that would be available at read time in a real storage scenario, i.e., no knowledge of the encoded DNA sequences other than the information received from sequencing.

The error analysis in **Figure *3b*** reveals an average error rate per position of 0.6%. Substitutions are the most prominent type (0.4%), twice as likely as deletions (0.2%) and ten times as common as insertions (0.04%). In some files, specific positions show higher error rates due to systematic errors in the reading or writing processes. Also, primer target regions (first and last 20 positions) suffer from fewer errors due to the nature of PCR, which favors amplification of perfect primers and primer target regions. A clear exception is the spike at position 9, which was caused by a single primer sequence with an error at that position. However, in all cases, errors in the primer region are low enough to associate most reads coming from sequencing to the sequenced files via sequence alignment. As such, we turn our attention to analysis of the non-primer region. **Figure *3c*** shows the percent breakdown of errors per base type, highlighting that insertion and substitution errors are biased toward certain base types. **Figure *3d*** also shows variation of error rates across differing neighboring base types.

Next, we proceeded to decode the data. In practice, current preparation and sequencing technologies yield some unusable reads due to their length being outside the acceptable range or too low confidence in base calling (5.5% on average in our experiments). We expect this number to improve as sequencing technology and wet lab protocols mature. We randomly subsampled the usable sequencing reads for each individual file, gradually increasing the number of reads supplied to the decoder. We were able to recover all 200MB of data (zero-byte difference when compared to original digital data) stored in the DNA with median coverage of only 5 reads per DNA sequence, with different files ranging from 3 to 14 reads per DNA sequence. If we include unusable reads in the calculation, the median goes up to 6.24 reads per DNA sequence. This is significantly lower coverage than was possible with earlier coding schemes (**Figure *1d***). The impact is lower cost because decoding from lower coverages allows a larger number of different DNA sequences to be read with the same sequencing kit. To understand the effect of coverage on our ability to decode files with no bit errors, we supplied the decoder with increasing coverage of reads, and measured the “Utilized Error Resilience” for several of our files (**Figure *3e***). As expected, the ratio decreases with coverage because the total number of errors and erasures decreases with extra read information.

We were also interested in understanding the impact of storage conditions on degradation. To assess this, we sampled two of our files, prepared them for sequencing, and divided them into three groups: (1) preserved by freezing at −20°C (~10% relative humidity [RH]), (2) dehydrated and kept at 65°C (~16% RH), and (3) treated with Biomatrica DNAstable and kept at 65°C (~16% RH). We aged them for 71 days and periodically sequenced the samples. Our evaluation metrics for degradation include (1) whether we can recover the file, and (2) how many of the original DNA sequences were missing in the sequencing data. All files were recoverable at all aging time points from samples originally containing the same amount of product with no additional preparation or material used. However, when compared to identical samples held at −20°C, the sample stored dry at 65°C had a larger set of unrecovered DNA sequences: 7,331 missing sequences versus 1,641 from the total set of 94,144 sequences (7.8% versus 1.7%). All sequences not recovered in the - 20°C sample were also not recovered in the 65°C sample due to their low initial copy number. The sample treated with DNAstable held at 65°C behaved similarly to the sample held at −20°C (1,691 vs 1,641, with 89 sequences missing from one but not both samples).

To further stress test our decoding scheme, we sequenced a 3KB file using the Oxford Nanopore Technologies (ONT) MinION sequencer. The compactness and potential for scalability makes nanopore-based sequencing an intriguing option for integration in future stand-alone DNA data storage systems. A key advantage is a very long read length of potentially thousands of nucleotides; however, with current technology, only a limited number of reads can be obtained from a single sequencing flow cell. To best utilize these characteristics, we developed a protocol to concatenate multiple oligonucleotides into longer reads. Using this approach, we successfully recovered the 3KB file sequenced with nanopore technology, despite a high coordinate error rate of 11.4%, computed using exhaustive minimum edit distance.

Given the current trends in data production and the rapid pace of progress of DNA manipulation technologies, we believe it to be possible to eventually replace tape, the densest commercially available storage media for archival storage. We contributed to a whole-system vision for DNA data storage by demonstrating a significant improvement in encoding and decoding, random access, and data volume stored in DNA (200MB, the largest synthetic DNA pool available to date). To encourage more work in this area, we will be making a limited number of samples of this pool available to select groups interested in DNA data storage.

## Methods

### Selectively amplifying DNA

First, the dehydrated single-stranded DNA pools obtained from synthesis were rehydrated with 1X TE buffer. To selectively amplify specific files, emulsion PCR (ePCR) was used due to its greater fidelity and ability to amplify sequences with less bias than PCR. We used the ePCR kit and protocol provided by CHIMERx. See supplementary information for more details.

The reaction was then put in a thermocycler with the following protocol: (1) 95°C for 3 min, (2) 98°C for 20 sec, (3) 62°C for 20 sec, (4) 72°C for 15 sec, (5) go to step 2 a varying number of times depending on the proportion of the pool being amplified, (6) 72°C for 30 sec. Note that we had a 25 random nucleotide sequence overhang on our forward primers, with the goal of introducing enough diversity to the first 25 cycles of sequencing to allow Illumina's NextSeq sequencing by synthesis cluster detection algorithm to identify clusters and pass quality filters. Interestingly, this is normally not necessary for genomic DNA since diversity is typical. After ePCR, the length of the dsDNA products was confirmed with a Qiaxcel fragment analyzer, with sample concentration measured by Qubit 3.0 fluorometer.

### Ligation of amplified DNA files for sequencing

After ePCR, amplified products were ligated to the Illumina sequencing adaptors with a modified version of Illumina TruSeq Nano ligation protocol and TruSeq ChIP Sample Preparation protocol. Briefly, samples were first converted to blunt ends, then purified with AMPure XP beads. An 'A' nucleotide was added to the 3’ends of the blunt DNA fragments, followed by ligation to the Illumina sequencing adaptors. We then cleaned the samples with Illumina sample purification beads and enriched the sample using PCR to yield enough product for sequencing. The length of enriched products was confirmed using a Qiaxcel bioanalyzer.

### Sample preparation for sequencing

When multiple separate samples were prepared for sequencing, these samples were mixed proportionally (e.g., a 10,000 oligonucleotide file to be sequenced with a 500,000 file would comprise 1.96% of the DNA material in this mix). The mixed sample was then prepared for sequencing by following the NextSeq System Denature and Dilute Libraries Guide. The sequencing sample was loaded into the sequencer at 1.3 pM, with a 10–20+% PhiX spike in as acontrol (PhiX is a reliable, adapter-ligated, well-characterized genomic DNA sample provided by Illumina).

### DNA degradation

Two files (with 66,656 and 27,488 sequences) were mixed, amplified, and fully prepared for sequencing. We aged the fully prepped libraries for 71 days and sequenced samples at intervals 0, 9, 31, 44 and 71 days. Three distinct sample preparation methods were compared: dry at 65°C, preserved in DNAstable at 65°C, and frozen in 15uL of elution buffer at −20°C. We successfully recovered both files with no additional effort from all samples at all time points.

### Sequencing with Oxford Nanopore Technologies MinlON

First, we used PCR to amplify an oligonucleotide library with primers containing orthogonal overhang sequences. Then, we combined the amplified products into one Gibson assembly reaction where each overhang allowed for multiple library members to be concatenated. Finally, we used PCR to amplify the resulting concatenated product with primers that hybridize to each respective end.

After dehydration, a 100-fold diluted sample of ssDNA library was amplified using a KAPA SYBR FAST qPCR kit with the following thermal profile: (1) 95°C for 3 min, (2) 98°C for 20 sec, (3) 69°C for 20 sec, (4) 72°C for 20 sec. The total number of cycles was determined by monitoring the fluorescence of the qPCR instrument as the amplification reached the plateau phase.

Three different amplification reactions were performed with primers containing distinct overhang regions necessary for a subsequent Gibson assembly reaction. Overhang sequences were designed using NUPACK ^14^ design module to avoid secondary structure formation. After amplification, each reaction was purified using Agencourt AMPure XP. Subsequently, amplification products mixed at equal molar ratio were added to NEB Gibson assembly master mix (1:1 volume ratio) and incubated at 50°C for 30 min.

Upon AMPure XP clean-up, the ligated product was amplified using the same qPCR protocol described above. Amplification was performed with primers corresponding to unique overhang sequences present at the 5’ and 3’ ends of the DNA. After amplification, a DNA band corresponding to the expected size was gel-extracted from a 2% agarose gel and quantified by a Qubit 3.0 fluorometer. The final product had a size of 590 bp including three fragments of the oligonucleotide library and the corresponding overhangs.

Sequencing sample preparation was performed according the Oxford Nanopore Technologies (ONT) Amplicon (R9) protocol. Metrichor sequencing metrics indicated 37,478 reads with workflow successful exit status out of 130,573 total reads. These reads were then successfully decoded the original digital file, despite the high coordinate error rate of 7.1%.

## Author contributions

L.O., Y.C., R.L. designed protocols and performed experiments. S.Y., S.D.A., K.M., M.R., C.R., P.G. designed and implemented the encoding and decoding pipeline. S.D.A., M.R., G.K., K.S., C.T., collected and analyzed data. B.N., C.T., S.N., G.G., H.P., R.C., and J.M. assisted in designing and evaluating experiments. D.C., G.S., L.C., and K.S. designed experiments, analyzed data and supervised the work.

P.G. is now at VMWare, Inc but contributed to this work when employed by Microsoft. G.K. is a graduate student at Stanford University but contributed to this work during an internship at Microsoft.

## Acknowledgements

We would like to thank Bill Peck, Patrick Finn, Siyuan Chen, Andrew Stewart, Bernadette Arias, and Emily Leproust, from Twist Bioscience for providing the DNA, suggesting protocol refinements and offering input to our data analysis. We also thank James Bornholt, Krittika D’Silva, and Anselm Levskaya for their help in the early stages of this project. This work was supported in part by a sponsored research agreement by Microsoft, NSF award CCF-1409831 to LC and GS and by NSF award CCF-1317653 to GS.

